# MORE IS DIFFERENT: DRUG PROPERTY ANALYSIS ON CELLULAR HIGH-CONTENT IMAGES USING DEEP LEARNING

**DOI:** 10.1101/2023.04.10.536183

**Authors:** Xiangrui Gao, Xueyu Guo, Fan Zhang, Mengcheng Yao, Xiaoxiao Wang, Dong Chen, Xiaodong Wang, Lipeng Lai

## Abstract

High-content analysis (HCA) holds enormous potential for drug discovery and research, but widely used methods can be cumbersome and yield inaccurate results. Noise and high similarity in cell images impede the accuracy of deep learning-based image analysis. To address these issues, we introduce More Is Different (MID), a novel HCA method that combines cellular experiments, image processing, and deep learning modeling. MID effectively combines the convolutional neural network and Transformer to encode high-content images, effectively filtering out noisy signals and characterizing cell phenotypes with high precision. In comparative tests on drug-induced cardiotoxicity and mitochondrial toxicity classification, as well as compound classification, MID outperformed both DeepProfiler and CellProfiler, which are two highly recognized methods in HCA. We believe that our results demonstrate the utility and versatility of MID and anticipate its widespread adoption in HCA for advancing drug development and disease research.

## INTRODUCTION

Developing novel pharmaceutical drugs represents a substantial investment that involves significant amount of time and resources, but with a low success rate. The major obstacles hindering drug development include ineffective drug activity, intractable drug toxicity, as well as marketing difficulties. However, the recent development of cellular phenotyping technology in drug discovery has demonstrated as a valuable tool to overcome the above issues. For example, Recursion’s phenotypic drug development system, the Recursion OS, has successfully driven several drugs to the clinical stage, showcasing its effectiveness ^[1]^.

Image processing and feature extraction are essential steps in HCA. One of the main tools currently in use is CellProfiler ^[2]^, developed by the imaging platform of Broad Institute, which relies on traditional image processing techniques. It enables biologists with no training in computer vision or programming to automatically identify and quantify phenotypes from thousands of images. Furthermore, deep learning models are increasingly employed for image feature extraction. Cimini and Carpenter et al. utilized ImageNet pretrained models for feature extraction and compared them with CellProfiler’s features. Their results showed some level of improvement, leading them to present DeepProfiler ^[3]^. This tool extracts cell-slice images based on the cellular localization provided by CellProfiler and uses a weakly supervised approach to classify DMSO and drug-treated images.

Today, the success of deep learning heavily relies on the availability of vast amounts of data and large advanced models. Massive data helps to train the model effectively without premature overfitting, while the reasonableness and complexity of the network structure endow the model with good memory and information extraction ability. In addition, the diversity of images within the enormous data makes training and prediction of models easier by eliminating complex preprocessing steps, such as segmentation, tracking, tracing, and spatial conversion. As a result, the task can perform end-to-end prediction directly, thereby simplifying the entire process.

However, simply migrating advanced computer vision models to HCA is not always successful, and larger models do not necessarily yield better results. For example, DeepProfiler, in practice, is less effective than a machine learning model based on cell phenotype features extracted by CellProfiler when predicting the drug properties of a compound. The problem lies in the fact that DeepProfiler is not optimized for cell images and only classifies individual cell image, making it difficult to be utilized for compound classification or compound property prediction from the cell-slice images ^[3]^. Moreover, a single-cell phenotype may not always represent the property distinction of a compound, as it may not have been influenced by the compound or may have died naturally during the experiment. Determining the phenotypic changes involving many cells provides a more reasonable resource for predicting a compound’s properties. Although CellProfiler counts the characteristics of many cells to make judgments, the phenotype of most cells may be undifferentiated, and some important information may be ignored during the counting process.

To address the aforementioned issues, we have developed MID - a deep learning-based method for processing cell phenotype images (Figure 1). The concept behind MID is inspired by P.W. Anderson’s quote “More is different” ^[4]^. In condensed matter physics, the behavior of large and complex elementary particle aggregates cannot be understood by simply extrapolating the properties of a few elementary particles. This also holds true for the problem of cellular phenotypes. During cellular experiments, the compound effects on cells are random, and the properties of compounds cannot be accurately distinguished from images of one or a few cells. Therefore, a more reasonable approach would be to consider many cell-slice images. By examining more cell-slice images, the model can learn to distinguish between important and unimportant images. Additionally, the information contained in multiple cell-slice images, even some contain noise, may reveal certain indiscernible patterns. Using a combination of convolutional neural networks and Transformers, MID can adaptively filter out irrelevant cell-slice images and extract features that accurately model cell phenotype characteristics. For different cell-slice images exposed to the same compound or compounds with similar drug properties, MID can provide similar features, while features extracted by MID differ more between compounds with distinct drug properties.

**Figure 1:**
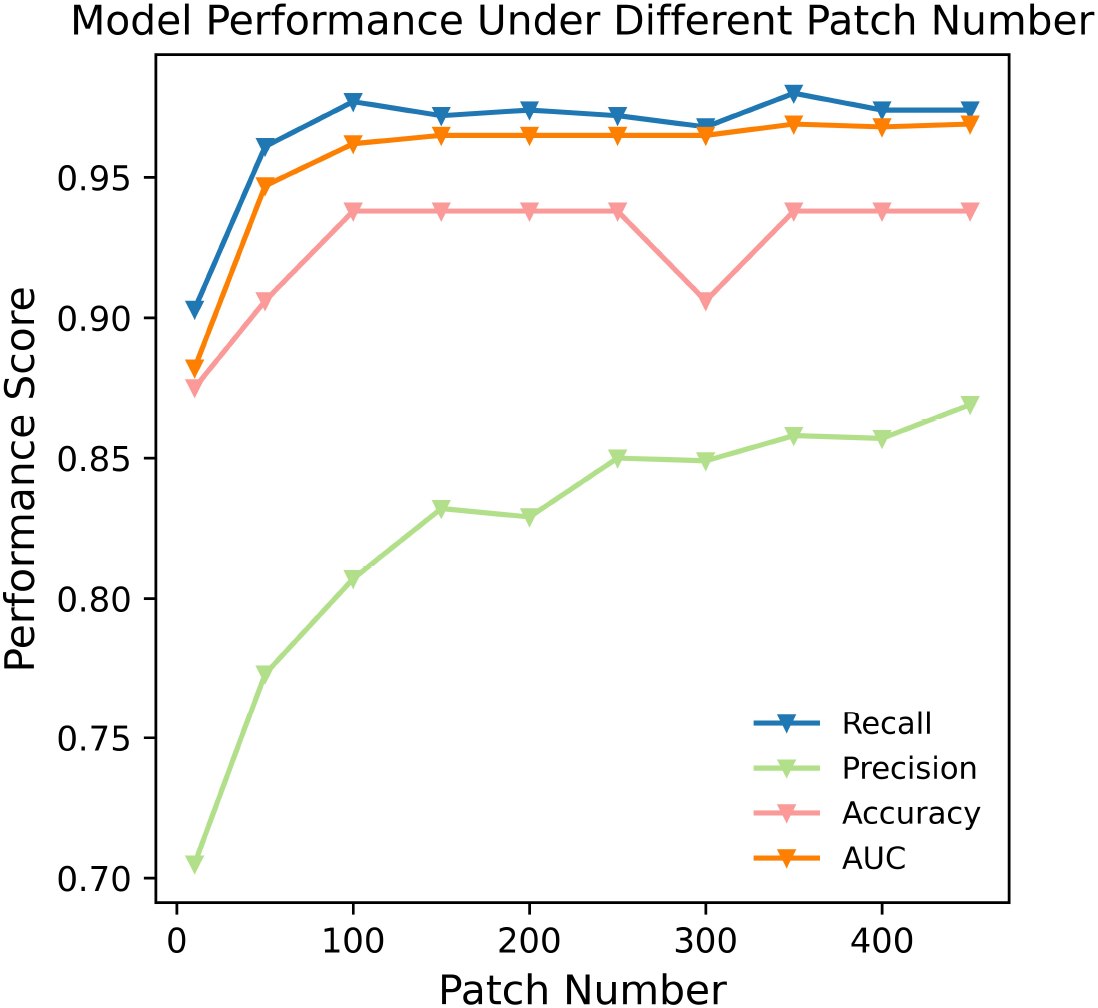
The performance of the MID (More Is Different) model under different patch number. The model prediction performance under different patch number of single cell-slice images in the test phase. The overall performance of the model increases with the increase of patch number.

Compared to DeepProfiler and CellProfiler, MID is a novel HCA process that leverages our understanding of cellular images and deep learning. It excels in screening and utilizing informative and high-quality cell-slice images to extract task-relevant cellular features, resulting in superior performance in three drug property-related tasks, i.e. 1) determination of drug inhibition on hEGR ion channels, 2) prediction of drug-induced mitochondrial toxicity, and 3) classification of compounds. In addition, MID can concatenate different compounds based on similar cell phenotypes, highlighting its potential in the field of drug repurposing and indication expansions.

## RESULTS

### MID performance on hERG inhibition classification

hERG, also known as Kv11.1, is a gene (KCNH2) that codes for an alpha subunit protein of the potassium ion channel ^[5]^. This channel, which is essential for the electrical activity of the heart, mediates the repolarizing IKr current in the cardiac action potential and coordinates heartbeats ^[6]^. However, some drugs in the market that inhibit hERG have the potential to prolong the QT interval and cause a dangerous irregularity of the heartbeat called torsades de pointes, making hERG inhibition an important factor to consider during drug development ^[7,8,9]^. Currently, there are several in vitro methods for evaluating hERG inhibition, electrophysiological (membrane clamp, the gold standard) ^[10]^, cytofluorimetric-based assays ^[11]^, and ligand binding assays ^[12,13]^, but they are often complicated, low throughput, and costly. We believe that the MID model can effectively determine whether a drug has the potential to inhibit hERG.

To test this hypothesis, we conducted high-content screening (HCS) experiments on 100 compounds, including water and DMSO, 57 of which were toxic and 45 non-toxic, using human induced pluripotent stem cell-derived cardiomyocytes (hiPSC-CMs). From these compounds, we selected 38 (19 non-toxic and 19 toxic) for the test dataset. We then used the images generated from our HCS experiments to evaluate the ability of three models - MID, DeepProfiler, and CellProfiler - to classify the cardiotoxicity of the drugs based on their ability to inhibit hERG channels.

Our results show that MID outperformed both DeepProfiler and CellProfiler, achieving an accuracy score of 90.6 in classifying images produced by drugs with or without hERG inhibition. We also calculated precision and recall scores for each model and found that MID had a precision score of 80.0 and a recall score of 95.5, indicating that it was capable of making correct predictions (Fig. 2a). Although CellProfiler-LightGBM had a higher recall score, it had a lower precision score, incorrectly predicting almost all negative cases as positive. In summary, on the classification task of hERG channel inhibition, MID performed best, followed by DeepProfiler, with CellProfiler performing the worst. Moreover, in this task, we try to demonstrate the differential impact of hERG channel inhibitors and non-hERG channel inhibitors on cell phenotype, as well as the effectiveness of MID in capturing this kind of difference. To achieve this, we randomly selected 10 compounds (5 labeled as 1 and 5 labeled as 0) and investigated their effects on cells at varying drug concentrations using MID model prediction scores. As depicted in Fig. 2d, the model evaluation scores exhibited distinct patterns for hERG channel inhibitors and non-hERG channel inhibitors as drug concentration increased. The scores for hERG channel inhibitors were higher, and with increasing concentration, the model evaluation scores increased and gradually reached a plateau or remained at a high level. The scores for non-hERG channel inhibitors were relatively low, and with increasing concentration, the model evaluation scores showed a trend of first increasing and then decreasing (possibly because these drugs have less impact on cells at low concentrations) or remained at a low level.

**Figure 2:**
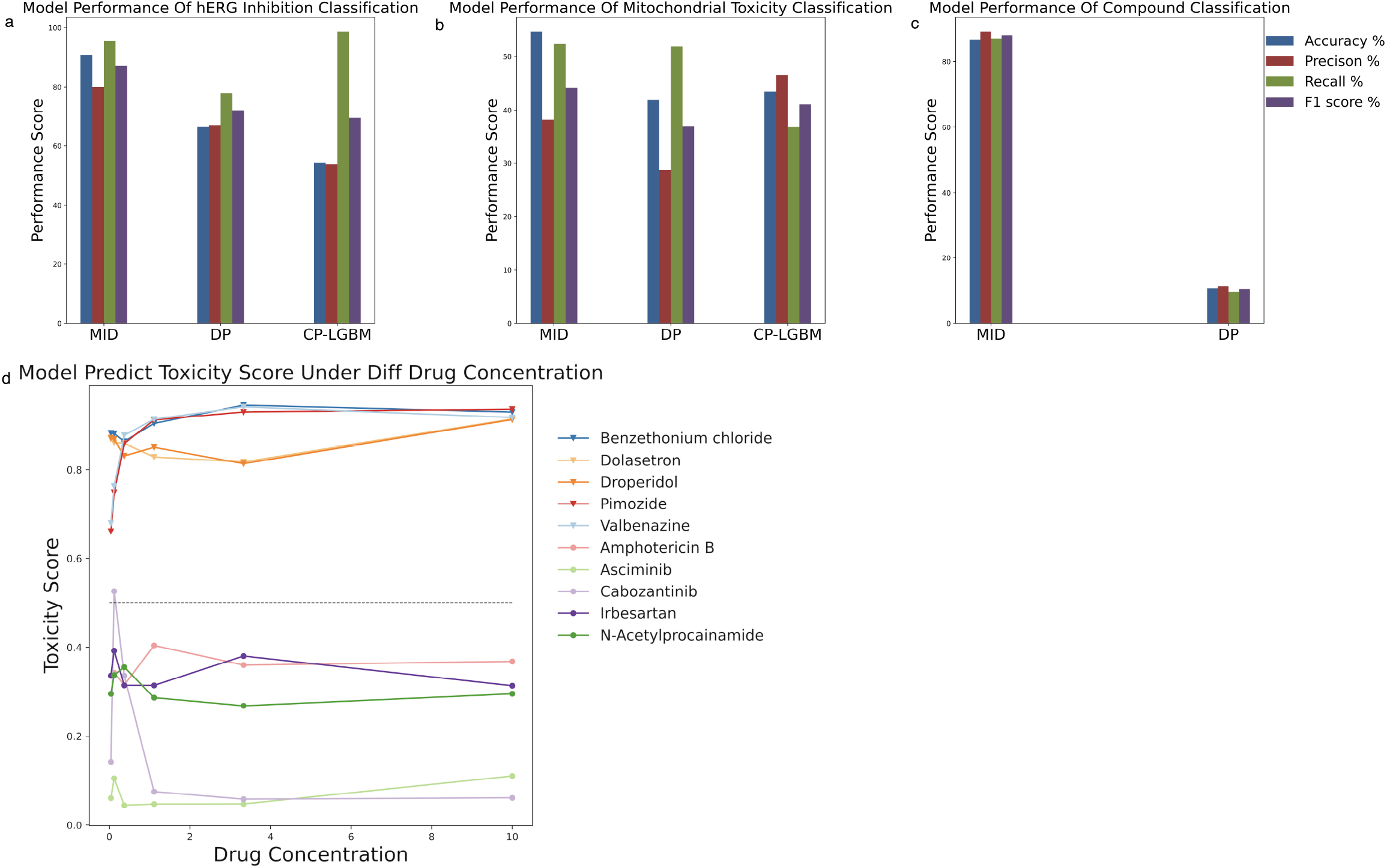
The performance of the three models on three different tasks. **(a)** In the hERG inhibition classification task, MID achieved the highest performance, DeepProfiler showed moderate performance, and the machine learning model using CellProfiler features performed the worst. **(b)** For the mitochondrial toxicity classification task, all three models performed consistently and comparably, with MID showing slightly better performance than the other two models in terms of specific metrics. **(c)** In the compound classification task, the MID model outperformed DeepProfiler. It’s worth noting that the machine learning model using CellProfiler features is not available in this table. **(d)** In hERG inhibiton classification task, the model evaluation scores exhibited distinct patterns for hERG channel inhibitors (shape in lower triangle) and non-hERG channel inhibitors (shape in dot) as drug concentration increased. The black dashed line represents y=0.5. Abbreviations used are DP for DeepProfiler and CP-LGBM for CellProfiler-LightGBM.

To fully interpret the performance of the MID model, we analyzed samples of its false predictions, including eltrombopag, pitolisant, and sildenafil, in the last section of the Results (MID bad cases contain information that deserves in-depth analysis).

### MID performance on Mitochondrial toxicity classification

Mitochondrial toxicity caused by certain compounds is a widespread form of organ toxicity, which can result in multiorgan damage in the heart, liver, bone, and brain ^[14, 15, 16]^. Regulators have withdrawn several drugs, including benfluorex, isoprenaline, nifedipine, and rosiglitazone, from the market due to their side effects, including mitochondrial toxicity and other adverse reactions ^[17]^. Furthermore, mitochondrial toxicity may be associated with drug resistance to linezolid ^[18]^. Therefore, mitochondrial toxicity is a critical factor in determining the success of drug development. Multiple mechanisms contribute to mitochondrial toxicity ^[19]^, resulting in various changes in cell phenotype. The alterations in cell morphology, texture, and intensity caused by compounds are strongly correlated with mitochondrial toxicity, suggesting that cell phenotype analysis is a reliable method for predicting mitochondrial toxicity[21].

In this study, we utilized public HCA images of 285 compounds from previous research ^[22]^. After labeling with PubChem annotation data, we selected a test set of 95 compounds, of which 18 were labeled ‘Active’, 51 were labeled ‘Inactive’, and 26 were labeled ‘Inconclusive’. Similar to the hERG task, we evaluated the classification performance of the three models. The MID model still outperforms the other two models in terms of specific metrics, as shown in Fig. 2b.

There may be two reasons for this. Firstly, the label’s definition is unclear. Secondly, the cellular portraits used in this task were stained with generic cellular dyes and were not specifically designed for observing drug mitochondrial toxicity. The use of general-purpose dyes to predict mitochondrial toxicity makes it more challenging for the models to capture the exact information or features that genuinely represent the mitochondrial toxicity of drugs, resulting in poor classification performance. In contrast, for the hERG task, we used cell dyes that specifically reflect the cardiotoxicity of drugs in HCA experiments. Customized experiments facilitated the MID model’s ability to capture information that is valid for making judgments. This demonstrates the importance of combining dry and wet experiments in areas related to drug discovery.

### MID performance on Compound classification

The compound prediction task involves classifying cell images based on their response to different compounds.In DeepProfiler, compound classification is a supervised training task that helps the model extract valuable image features. Our results showed that the MID model outperformed DeepProfiler, achieving an accuracy score of 86.7 compared to 10.7, a precision score of 89.2 compared to 11.4, a recall score of 87.1 compared to 9.7, and a F1 score of 88.1 compared to 10.5. Note that the machine learning model that utilizes CellProfiler features is not available here (Fig. 2c).

### MID can act as an effective cell-slice filter

To demonstrate the ability of MID to extract relevant information, we generated an attention heatmap using MID, which highlighted the correlation between cells in the HCA image and the cardiotoxicity task (Fig. 3a and Fig. 3b). As indicated by the self-attention mechanism, the similarity between the embeddings of single cell-slice image and the embeddings of the CLS token is positively associated with the classification task ^[23]^. In the heatmap, cells that are more relevant to the downstream task are represented by brighter and warmer dots, and the dots in Fig. 3b clearly illustrate the differences between different cells in terms of task correlation.

**Figure 3:**
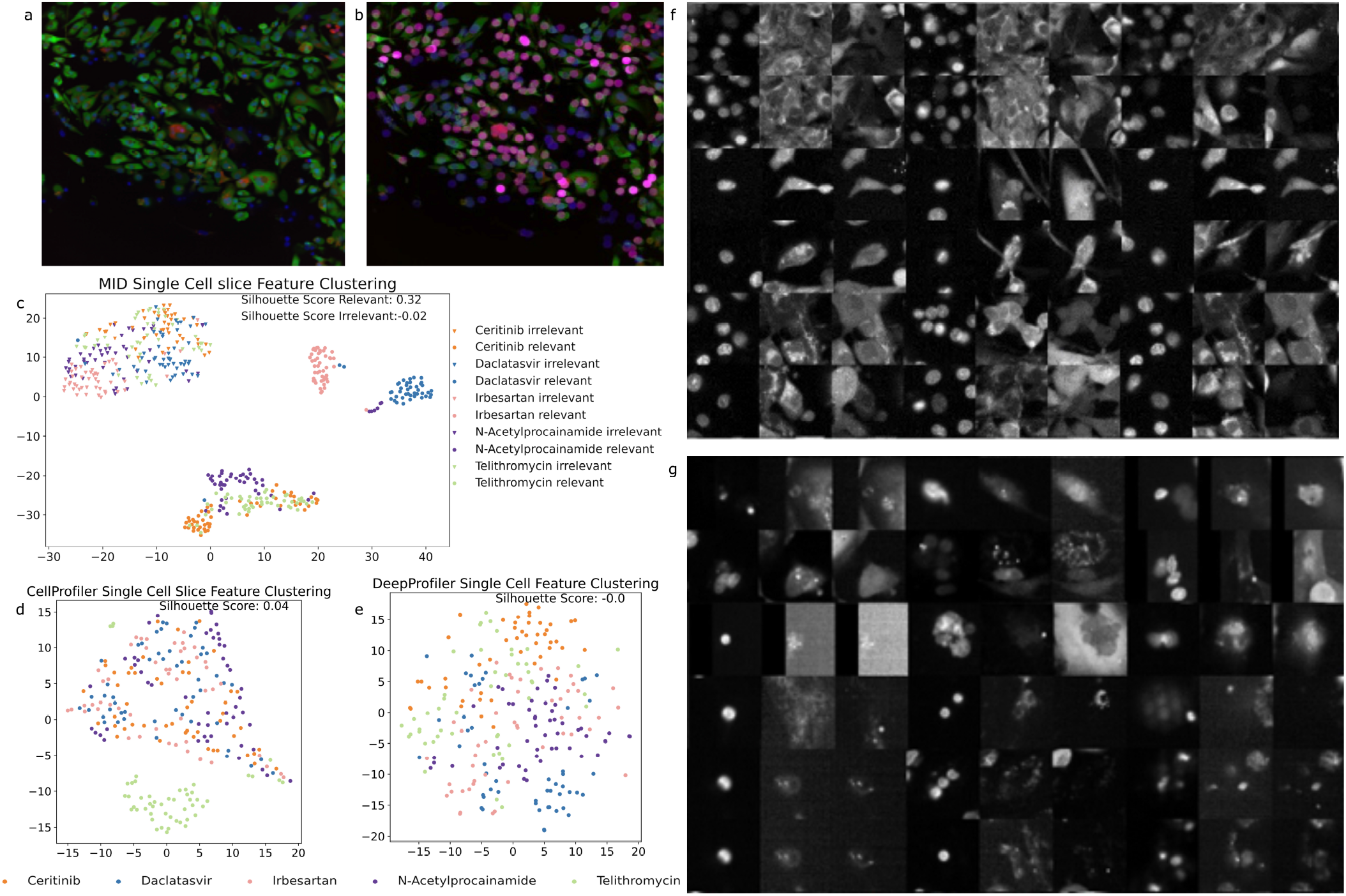
MID can act as a filter to identify cell slices that are relevant to a downstream classification task. We used an attention heatmap to visualize the correlation between the CLS embeddings of cell phenotype images and the downstream task. Cells that are more relevant to the task are highlighted in brighter, warmer colors. We added these colors to the Cell Painting image to demonstrate the ability of MID to recognize task-related and nonrelevant cells. By clustering the MID embeddings of relevant and irrelevant cell-slice images from different compounds, we found that MID can effectively filter out nonrelevant cells and identify those that are relevant for classification. The relevant images have a deeper staining level, more intact cell structure, and higher quality than the irrelevant images. (**a)** Cell painting image of cells treated with alfuzosin. Nucleus in blue, ROS in green, MMP in red. **(b)** Attention heatmap of the cell painting image of cells treated with alfuzosin. Some randomly selected cells in the cell painting image were marked with colored solid dots. The color changes gradually from dark blue to bright pink, which means that the marked cells are increasingly relevant to the downstream task. **(c)** We clustered the MID embeddings of relevant (shape in dot) and irrelevant (shape in lower triangle) cell-slice images of 5 compounds and found that the relevant cell-slice images from different compounds clustered separately and were apart from each other. In contrast, all the irrelevant cell-slice images gathered into a large cluster regardless of which compound they belonged to. This phenomenon indicates that the multiple cell slices classifier in our MID model can act as a cell filter. **(d)** Feature clusters of CellProfiler of cell-slice images. For each compound, we randomly selected 50 cell slices **(e)** Feature clusters of DeepProfiler of cell-slice images. For each compound, we randomly selected 50 cell slices. Compared with MID, the cell-slice feature distribution of CellProfiler and DeepProfiler was more dispersed; on the one hand, features of the same compound were not aggregated into a compact cluster, and on the other hand, features of different compounds were mixed. **(f)** Relevant images of belzutifan, carvedilol and daclatasvir. Each three columns (the first column is Nucleus, the second is ROS, the third is MMP) correspond to one compound and each compound has six rows of sampled images, of which the cell staining level is sufficient, the cell structure is clear and easy to read, and the image is of high quality and has less background noise. **(g)** Irrelevant images of belzutifan, carvedilol and daclatasvir. Each three columns (the first column is Nucleus, the second is ROS, the third is MMP) correspond to one compound and each compound has six rows of sampled images, of which the cell staining level is inadequate, the cell structure is ambiguous, and the quality is unqualified and full of noise. ROS, mitochondrial reactive oxygen species. MMP, mitochondrial membrane potential.

To determine whether the Transformer can isolate cell slices of interest, we analyzed the features of task-relevant single cell-slice image and the irrelevant ones, and based on the attention map values, we can define how relevant the image is to the task. As shown in Fig. 3c, using MID embeddings, the cell slices unrelated to the cardiotoxicity of the compound are tightly clustered regardless of whether the compounds are identical, while the cell slices related to drug-induced cardiotoxicity are separately clustered according to compounds, with most of the compounds widely spaced apart. This demonstrates that the multiple cell slices encoder can function as a cell filter. The clustering results also provide two other insights. Firstly, in cellular experiments using different compounds with different properties, there are always some dead cells or normal cells that are not affected by the compound, which is the common part and does not contribute much to classification. The characteristics of all these cells after the single cell-slice encoder are similar, and their clustering together indirectly confirms that the model has learned the correct information. Secondly, after being exposed to different compounds, the phenotype of the cells changes more significantly and they become very distinct from each other, even when they are dead. We also clustered the single cell-slice image features extracted by CellProfiler (Fig. 3d) and DeepProfiler (Fig. 3e), and because CellProfiler and DeepProfiler did not distinguish whether those single cell-slice images were task-related, the features they extracted were not representative of different compounds and had low discriminatory power compared to MID.

### Task-relevant cell-slice images are highly informative

To further demonstrate the screening capabilities of MID for cell phenotype images, we analyzed and compared cell-slice images that were considered relevant and irrelevant based on the MID attention map (Fig. 3f, Fig. 3g). Each cell-slice image was comprised of three channels: nucleus, mitochondrial reactive oxygen species (ROS), and mitochondrial membrane potential (MMP). Highly correlated images (Fig. 3f) exhibited deeper staining, higher fluorescence intensity, greater contrast between channels, and more accurate staining in the corresponding channels, as compared to the low-correlation images (Fig. 3g). Furthermore, highly correlated images displayed more distinct cell structure in the ROS and MMP channels, resulting in more accurate cell identification, while most cells in the low-correlation images exhibited the opposite characteristics. In terms of image quality, high-correlation images were clear and free of impurities, whereas low-correlation images had a halo on the image surface due to optical structure formation, leading to indistinct images and high noise. Thus, the attention mechanism utilized in MID allowed for the extraction of highly informative, accurate, and structurally intact cell-slice images while filtering out low-informative, erroneous, and poor-quality images, thereby improving the performance of the model after image preprocessing.

### Multiple cell slices enable accurate grasp of the compound property

To illustrate the core idea of MID, we clustered the features from single cell-slice relevant images and multiple cell slices relevant images (Figure 4). First, using the MID features of single cell-slice images (Fig. 4a), the clusters of different compounds were mixed compared to those of multiple cell-slice images (Fig. 4b). This suggests that incorporating more image slices provides more information, and MID can anchor the useful parts from numerous single cell-slice images and integrate them to predict the specific property of compounds. The feature clusters of compounds (benzethonium chloride and betrixaban) changed from being mixed (Fig. 4a) to forming independent clusters (Fig. 4b), which verifies this idea well. Secondly, when comparing the feature clusters of single cell-slice images obtained using MID with those obtained using CellProfiler (Fig. 4c) and DeepProfiler (Fig. 4e), the distances between the single cell-slice image embeddings of MID, under the influence of different compounds, were still large, and their boundaries were still distinct (Fig. 4a). Moreover, the feature clusters of MID using multiple cell-slice images (Fig. 4b) were more accurate compared to those of CellProfiler (Fig. 4d) and DeepProfiler (Fig. 4f). When using multiple cell-slice images, the advantages of MID are evident as it can efficiently differentiate between different compounds. The clustering of belzutifan, carvedilol and daclatasvir, which are closely clustered compared to other compounds (Fig. 4b), may suggest that the cell phenotype images corresponding to these three compounds are similar.

**Figure 4:**
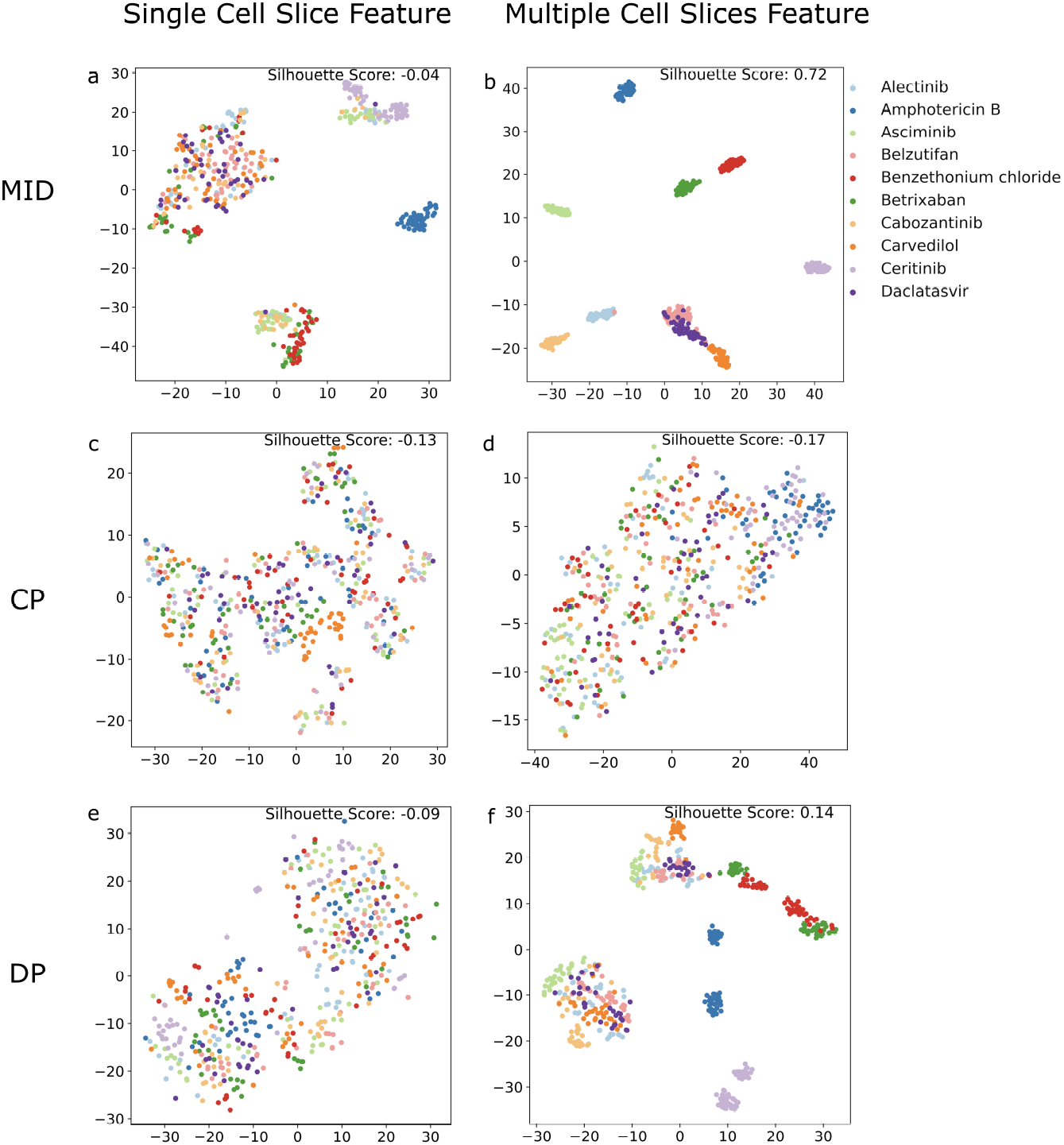
Feature clusters of single cell-slice image and multiple cell-slice images. **(a)** The feature clusters of single cell-slice images using the MID model. Similar to CellProfile **(c)** and DeepProfiler **(e)**, the feature distribution of the 10 compounds appeared scattered and confounded, making it impossible to distinguish between different compounds. **(b)** The feature clusters of multiple cell-slice images using the MID model. The MID approach captures enough relevant information from multiple cell-slice images to be representative of the sample. Clustering using features of multiple cell-slice images showed that the same compounds were clustered together and clusters of different compounds were dispersed at long distances, indicating that these features have a high degree of discrimination. **(d)** The feature clusters of multiple cell-slice images using CellProfiler, which uses simple averaging to fuse information from multiple cell slices to represent the sample. However, this simple mechanical integration employed by CellProfiler was insufficient to obtain information related to the properties of the compound when compared with MID. **(f)** The feature clusters of multiple cell-slice images using DeepProfiler. Although the effect of DeepProfiler was better than that of CellProfiler, it still lagged behind the performance of MID. Furthermore, when using the features of multiple cell-slice images, the clusters of compounds benzethonium chloride and betrixaban were distributed very separately, and the individual cluster was more compact, indicating the advantage of using multiple cell slices. That is, the MID method can extract valid information from all the useful cell-slice images to represent the tested sample. Abbreviations used are DP for DeepProfiler and CP for CellProfiler.

### The MID embeddings can represent the cell phenotypes

If the embedding of MID can accurately represent the cellular phenotype, this suggests that compounds whose MID embeddings cluster together are likely to have very similar cell phenotypes. To test this hypothesis, we compared images of three drugs (belzutifan, carvedilol, and daclatasvir) as shown in Fig. 5a. However, it was challenging for us to discern the differences between the three drugs based solely on their cellular phenotypes.

**Figure 5:**
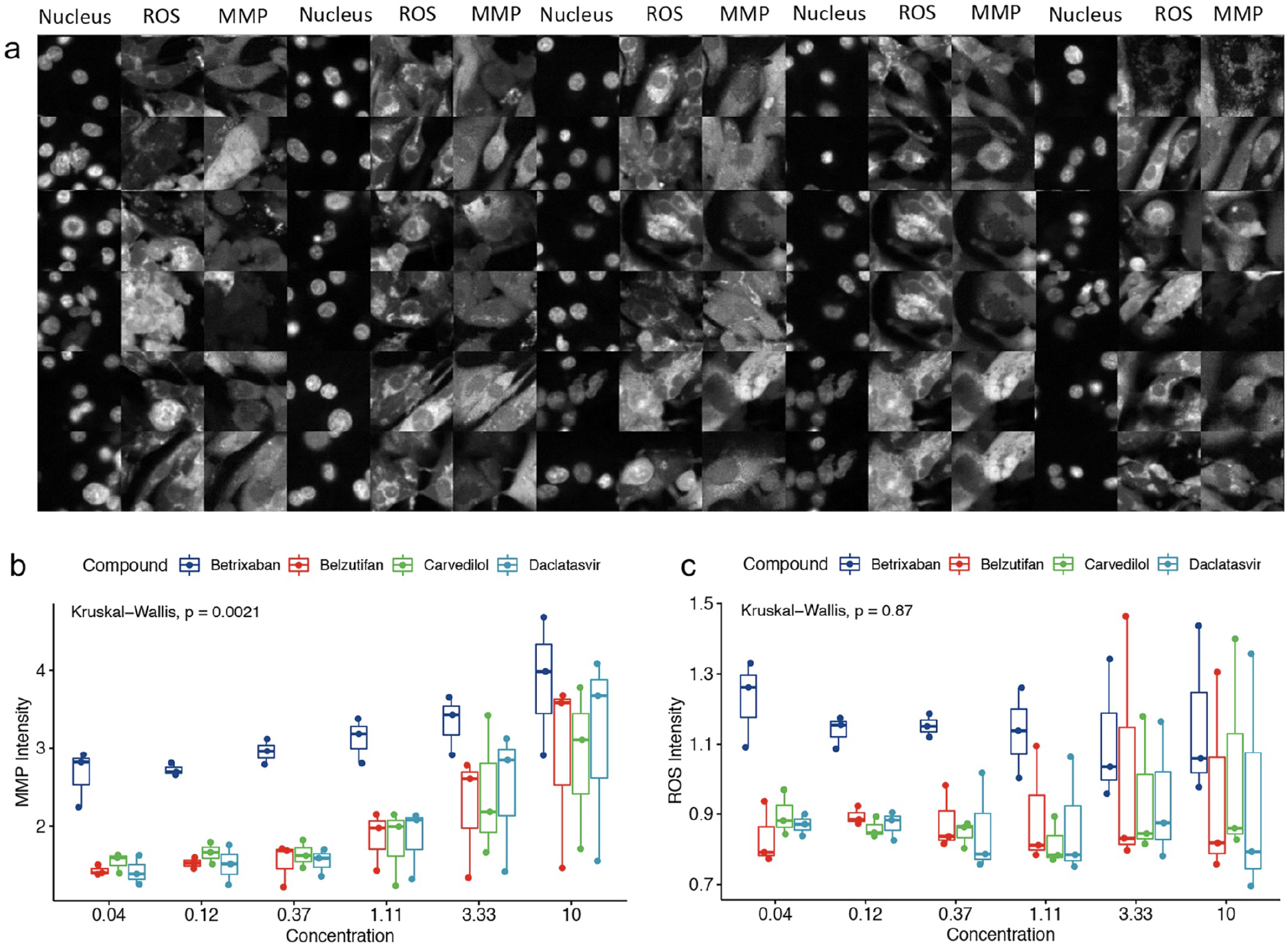
Analysis of cell phenotype similar compounds show that MID embeddings can represent the cell phenotypes. **(a)** cell-slice images of three compounds clustered together by MID embeddings in the hEGR classification task. For the three compounds (belzutifan, carvedilol and daclatasvir), each compound has two rows of images, where the first row is for relevant images and the second row is for irrelevant ones. The three compounds are clustered together in Fig. 4b, and we cannot differentiate any one of them from each other based on the images shown here, regardless of whether they are relevant or irrelevant, which illustrates the consistency of the cell phenotype images and the MID embeddings. ROS, mitochondrial reactive oxygen species. MMP, mitochondrial membrane potential. The intensity of MMP **(b)** and ROS **(c)** calculated from cell images separately perturbated by belzutifan, carvedilol, daclatasvir and betrixaban, of which the first three drugs clustered together and separated from the fourth drug in Fig. 4b. The MMP or ROS intensity of the first three at different concentrations shows a nearly consistent level while significantly differing from the fourth, which interprets the judgment of MID and the similarity of cell phenotype for the first three drugs. ROS, mitochondrial reactive oxygen species. MMP, mitochondrial membrane potential.

Then, we separately analyzed the changes in MMP and ROS intensity of cells treated with different concentrations of three drugs, each of which may have different effects on certain cell statistical indicators. We also compared the effects of these three drugs with that of betrixaban, a drug that is much farther away in terms of embedding distance. Our results show that the changes in MMP (Fig. 5b) and ROS (Fig. 5c) intensity are similar for the three drugs with similar cell phenotypes, while betrixaban behaves differently.

To better understand the results presented in Figure 5, we researched the mechanism of action (MOA) of the three drugs in question. Despite their different indications - Belzutifan for von Hippel-Lindau (VHL) syndrome-associated clear-cell renal cell carcinoma (ccRCC), Carvedilol for hypertension, and Daclatasvir for Chronic hepatitis C genotype 3 (GT-3 HCV) - their MOAs suggest that they may have similar effects on cardiomyocytes.

Belzutifan inhibits hypoxia-inducible factor 2α (HIF-2α), and research suggests that HIF-2α inhibitors can reverse pulmonary hypertension and that there are shared pathophysiologic mechanisms between cancer and heart failure ^[24, 25]^. Carvedilol, a nonselective beta-adrenergic antagonist, has anti-free radical and antioxidant effects, and can resist oxidation and reduce ROS production ^[26]^. It also inhibits the Cardiac Mitochondrial Permeability Transition (MPT), which can depolarize mitochondrial membranes and uncouple oxidative phosphorylation (OXPHOS) ^[27]^. Daclatasvir, a pangenotypic NS5A replication complex inhibitor with a dual antiviral effect, inhibits RNA replication and viral assembly. Cellular ROS levels rise during HCV infection, and evidence suggests that anti-RNA viral drugs are associated with intracellular ROS levels ^[28]^. ROS may also play a critical role as a signal molecule in the regulation of viral replication and organelle function ^[29]^.

### MID bad cases contain information that deserves in-depth analysis

During the hERG classification task, our model inaccurately identified three drugs, eltrombopag, pitolisant, and sildenafil, as cardiotoxic based on hERG IC50 values, but being predicted as non-toxic by MID. To understand the reasons behind these false predictions, we analyzed and compared the cell painting staining images of cells treated with toxic and non-toxic compounds.

Figure 6 displays the cell painting staining results of the cells after treatment with the compounds, where the nucleus is blue, ROS is green, MMP is yellow, and dead cells are red. The images depicted in Fig. 6b represent the staining results of three significantly non-toxic compounds, namely irbesartan with hEGR IC50 values of 194 μM, riluzole with 25 μM, and belzutifan with 50 μM. The staining channels showed an intact cell structure with uniformly distributed green fluorescence and almost no dead cells. The images shown in Fig. 6c illustrate the staining results of three significantly toxic compounds, namely nintedanib with hEGR IC50 values of 2.6 μM, sunitinib with 0.25-10 μM, and ponatinib with 0.77 μM. In nintedanib, green fluorescence presented a punctate distribution around the nuclei, and dead cells were visible. In sunitinib, the number of dead cells increased, and most of the cells lost their intact cell structure. In ponatinib, almost all the cells were dead. Based on the recognizable cardiotoxicity represented by the images, for the compounds with incorrect predictions shown in Fig. 6a, the images were more similar to those in Fig. 6b. That is, the cells had an intact structure, the green fluorescence was uniformly lamellar, and there were almost no dead cells. Therefore, we suggested that the model tends to identify these three compounds as non-toxic. Additionally, as shown in Supplementary Figure 1, the clusters of MID embeddings of all 100 compounds, plus DMSO and water (N), eltrombopag was located close to irbesartan and surrounded by six other non-toxic compounds (darunavir, lesinurad, omarigliptin, miltefosine, glibenclamide, and rosiglitazone). Pitolisant was close to belzutifan, and sildenafil almost overlapped with riluzole. Furthermore, based on some previous literature, we found that none of the three drugs themselves had significant cardiotoxicity in clinical or nonclinical studies ^[30,31,32,33,34,35]^. In summary, the comparison of cell painting images and supporting evidence from the literature suggest that the results of MID’s wrong predictions are informative and worthy of attention.

**Figure 6:**
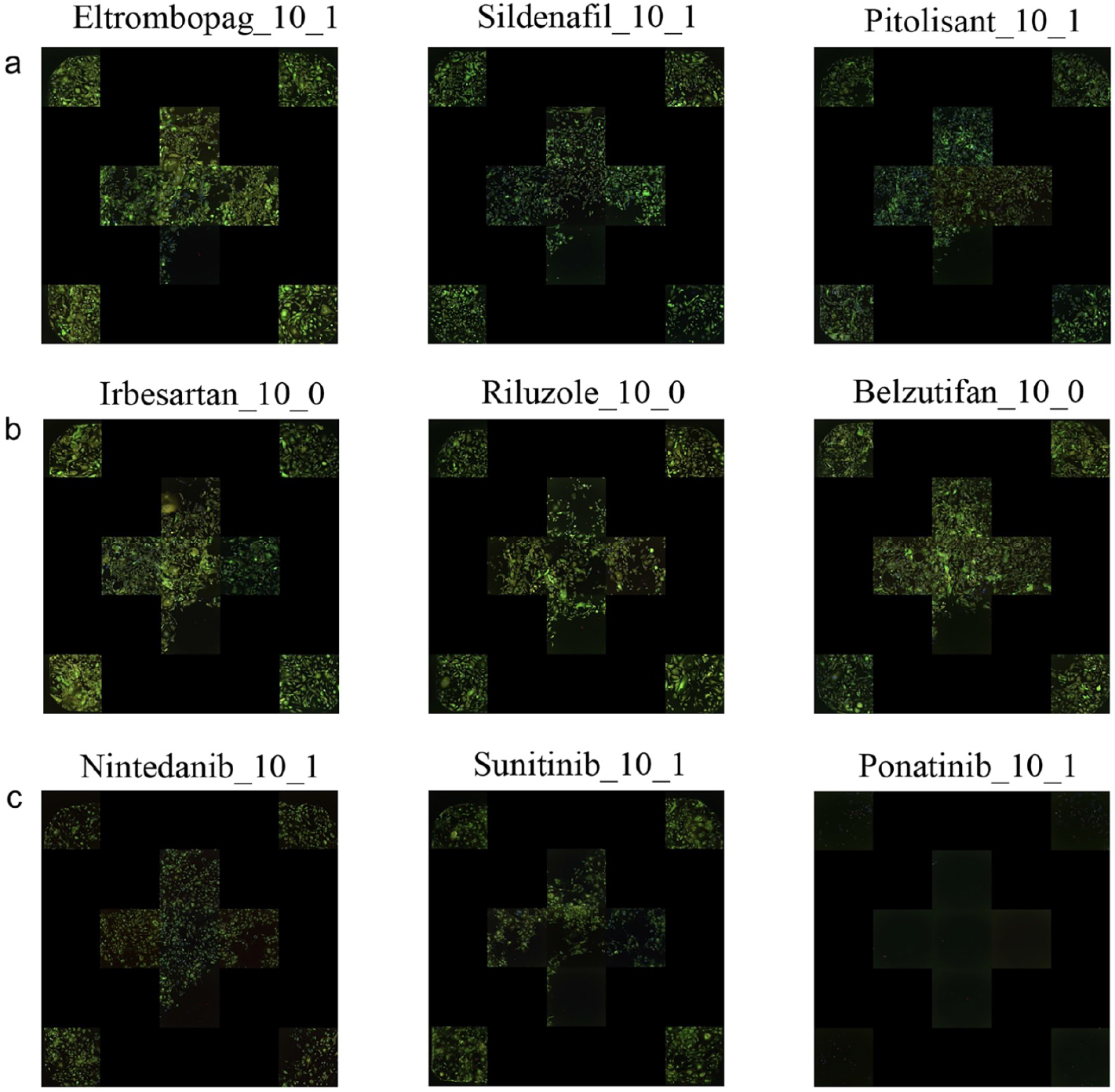
The results of cell painting staining for cells treated with compounds that either do or do not exhibit cardiotoxicity. **(a)** The staining results for three compounds that MID model incorrectly predicted as nontoxic. **(b)** Cell Paintings staining results of cells treated with three significantly nontoxic compounds, in which the cell structure is intact with uniformly distributed green fluorescence and nearly no dead cells. **(c)** Cell painting staining results of cells treated with three significantly toxic compounds, in which the cell structure is cracked with separately distributed green fluorescence and dead cells are clearly visible. Notably, the staining patterns for the incorrectly predicted compounds in (a) are more similar to those observed for nontoxic compounds.

## DISCUSSION

In this study, we developed a highly accurate computational model (MID) using a deep learning framework to extract relevant information from high-content images in HCS experiments for drug property analysis. We thoroughly studied the characteristics of cell phenotype images to construct the method from data processing to model training and validation. To assess the potential value of our method in high-content cellular image processing, we selected three widely used downstream tasks in drug discovery: Task 1 - classification of drug inhibition of the hERG ion channel, Task 2 - classification of drug mitochondrial toxicity, and Task 3 - classification of compounds.

To validate our approach, we designed HCS experiments using a batch of compounds with known hERG IC50 values to produce cell images for tasks 1 and 3. We used open-source high-content image data to construct datasets for task 2. After applying MID to the downstream tasks, we analyzed the results in detail. Our findings indicate that MID outperforms CellProfiler and DeepProfiler in tasks 1 and 3. Additionally, MID performs better than CellProfiler and DeepProfiler in task 2, with results consistent with those reported in the literature ^[21]^ (Figure 2).

In the hERG task, we compared the original cell-slice images input to the MID model and analyzed the valid and invalid information extracted by MID. Our analysis revealed that MID accurately identifies informative and high-quality cell-slice images using a self-attention mechanism (Figure 3). We also found that MID predicts the properties of compounds more accurately when using more cell-slice images (Figure 4). Furthermore, we analyzed biologically significant cell phenotype features extracted by CellProfiler for images judged to be similar by MID but disturbed by different drugs. We found that compounds with different indications but similar cell phenotypes, such as belzutifan, carvedilol, and daclatasvir, were highly consistent in the ROS and MMP intensities of the corresponding images (Figure 5). This confirms the accuracy of the judgment from MID and suggests that MID models may be useful in high-throughput applications in the field of drug indication expansion and drug repurposing. Lastly, we analyzed MID bad cases by comparing images and searching the literature. Our analysis suggested that three compounds defined as cardiotoxic by hERG IC50 values were nontoxic according to the image, literature, and MID predictions (Figure 6). Overall, our results demonstrate that MID is a highly effective tool for high-content cell image analysis, with potential applications in drug discovery and development. In conclusion, HCS is an emerging field that is still rapidly evolving in terms of experimental implementation and analytical methods and has the potential to solve diverse biological problems. The key to achieving credible results in downstream tasks lies in the ability of analytical tools or computational models to select useful parts from a large number of cell images of varying quality. Among the tested solutions for downstream tasks, CellProfiler and DeepProfiler performed mediocre, while MID showed promising results. This is because MID, which leverages deep learning to use multiple cell slices simultaneously for model training and verification, can accurately capture key information and eliminate noise interference. Additionally, MID can distinguish between cells with similar phenotypes but perturbed by different compounds, suggesting many possibilities for downstream applications. Overall, the conclusion emphasizes the potential of MID as a valuable tool for high-content cell image analysis, which can contribute to various fields such as drug discovery and development, disease diagnosis, and personalized medicine.

In the next phase of our research, we plan to expand the application of MID to more cell phenotyping tasks and high-content images induced by other perturbations, such as RNA interference (RNAi) or CRISPR-Cas9. Our goal is to demonstrate the versatility and effectiveness of our model in analyzing various types of cellular image data. However, we also acknowledge that algorithms based on cell phenotype images have limitations, particularly when the changes in cell phenotype induced by a compound are minimal, which may make it difficult for the model to accurately assess its toxicity or other characteristics. To address this issue, we plan to explore the integration of other high-level information, such as transcriptome data or videos of cells under bright field, using multimodal learning techniques. We believe that incorporating multiple sources of information will enhance the accuracy and robustness of our model and enable it to handle a wider range of biological problems.

## METHODS

### Constructing the cardiotoxicity high-content image data

To obtain an appropriate image dataset for the hERG inhibition task, we selected 100 compounds for HCS experiments and generated corresponding high-content images using a high-content imager. Supplementary Table 2 provides detailed information for each compound, including plate name, drug name, hERG IC50 value range, and cardiotoxicity label. We first downloaded all compounds with hERG IC50 values using the python API interface of the chEMBL Database and subsequently selected 100 compounds by deduplicating and filtering drug molecules with ambiguous hERG IC50 values. The compounds were then labeled as cardiotoxic and noncardiotoxic based on their hERG IC50 values using a threshold of 10 μM (less than or equal to 10 μM for toxic labeling and greater than 10 μM for nontoxic labeling). This resulted in 47 cardiotoxic and 53 non-cardiotoxic compounds, on which we performed cellular experiments using hiPSC-CMs.

The hiPSC-CMs at day 30 after cardiac induction were cryopreserved as Cauliscell hiPSC-CMs (Cauliscell Company, Nanjing, China) and thawed in a 37°C water bath with gentle shaking. After centrifugation and counting, the cells were added to 384-well plates precoated with 10 μg/ml recombinant human vitronectin at 12,000 cells/well (Cauliscell) in cardiomyocyte plating medium (Cauliscell). After 24 hours, the volume was replaced with cardiomyocyte maintenance medium, which was changed every other day. Once the cells started to beat rhythmically, we added the compounds, which were selected from the FDA-approved compound library (FDA-approved drug library, MedChemExpress) for cardiotoxicity testing. The working concentrations of each compound were 10 μM, 3.33 μM, 1.11 μM, 0.37 μM, 0.12 μM, and 0.04 μM, with 3 replicate instances set for each compound at each concentration. The control group was set with 0.1% DMSO (Sigma), and the blank control was set with water. After incubating the cells with compounds for 72 hours, working concentration dyes of CM-H2DCFDA (Thermo) at 5 μM, TMRM (Thermo) at 20 nM, and Hoechst33342 (Thermo) at 5 μg/ml were prepared with maintenance medium (Cauliscell) and added to the cells for 30 min. The cells were washed with HBSS (Beyotime Biotechnology), and YOYO-3 (Thermo) was added at a final concentration of 2 nM, followed by incubation at room temperature for 10 min and washing with HBSS (Beyotime Biotechnology). Finally, the 384-well plate (PerkinElmer) was placed on a high-content imager (Operetta CLS, PerkinElmer), and images were taken with a 20x water objective in the confocal model with 9 fields of view for one well, corresponding to the filter parameters shown in Supplementary Table 1.

### Constructing the mitochondrial toxicity high-content image data

For the mitochondrial toxicity task, we utilized open-source high-content images from Bray et al. ^[20]^. A total of fifty-five plate high-content images with numerous compounds were downloaded, and the mitochondrial toxicity labels, obtained from the PubChem Database (https://pubchem.ncbi.nlm.nih.gov/assay/pcget.cgi?query=download&record_type=datatable&actvty=all&response_type=save&aid=720637), were assigned to each compound. After labeling, 285 compounds were collected, of which 47 were classified as active, 141 as inactive, and 97 as inconclusive. Detailed information regarding the drug names and mitochondrial toxicity labels from PubChem can be found in Supplementary Table 3. Supplementary Table 4 shows the datasets and models used in the three tasks.

### CellProfiler and DeepProfiler data processing

For the cardiotoxicity and mitochondrial toxicity high-content images, we extracted cell phenotype features using standard procedures of CellProfiler (version 4.2.4). Subsequently, we utilized the machine learning model called LGBMClassifier from scikit-learn to perform classification tasks on toxicity data, after dealing with cell phenotype features. Once the locations of single cells were identified in the images by CellProfiler, we inputted the images and locations to DeepProfiler (version 0.3.1). During classification tasks, we employed a pretrained model named EffientNet, which was deployed within DeepProfiler, and calculated embeddings using DeepProfiler. The CellProfiler-LightGBM and DeepProfiler results were evaluated and compared with MID to assess their accuracy and generalizability. In all three models (MID, DeepProfiler and CellProfiler-LightGBM), we partitioned the 285 compounds labeled with mitochondrial toxicity into a training set comprising 190 compounds and a test set containing 95 compounds. For the 100 cardiotoxic compounds, the training and testing sets contained 68 and 32 compounds, respectively.

### ROS and MMP Measurement

To measure ROS and MMP, we utilized cell phenotype features calculated from our high-content images of 100 compounds, which were used for cardiotoxicity assessment. We designed two CellProfiler analysis protocols that can automatically detect and quantify fluorescence intensity, which proved to be useful for analyzing large image datasets. The ROS and MMP measurements were obtained from the suppressed fluorescent channels of the H2DCFDA and TMRM cell-based assay kit. We computed a reduced dataset with the well-mean feature vector per well, followed by normalizing all features by subtracting the mean of each plate layout from each feature.

### MID model Design

Cell phenotype images are distinct from general images in two key ways. First, they contain a high degree of redundancy, with dark backgrounds and bright cells dominating most of the image. As a result, it can be challenging to distinguish differences between cells using generic training. Second, there is a lot of noise in cell phenotype images due to experimental manipulations and batches, requiring different concentration gradients and experimental replicates to eliminate stochastic factors. Existing models and processing techniques have struggled to address these challenges, resulting in poor results.

To address these issues, we developed the MID model, a deep learning model specifically designed for cell phenotype images. MID is a plug-and-play flexible model framework that is not restricted to a particular backbone. The model processes cell-slice images by normalizing and grouping them by compound and concentration. First, single cell locations are extracted from CellProfiler by calculating the center coordinate of the nucleus. Second, images are rescaled with the global fluorescence intensity, and third, DeepProfiler crops cell slices from 3 channels based on the x and y coordinates for the center of a single nucleus. In all experiments, cell slices were cropped from a region of 96 × 96 pixels centered on the nucleus without resizing. The resulting 96 × 288-pixel images were preserved for model construction. Cell-slice images stained with different dyes were integrated as different channels into a single cell-slice multichannel image. This approach reduces noise interference and allows the model to perform the classification task using a few representative cells. Furthermore, the training phase inputs a limited number of cell images at a time to prevent the model from overfitting prematurely, which brought by complex information of too many cell images.

The MID model consists of three main parts: a single cell-slice encoder, a multiple cell slices encoder, and a classifier. The single cell-slice encoder uses a uniform CNN network to obtain an unbiased latent representation of single cell-slice image and improve the generalization of the latent representation. The multiple cell slices encoder is built by Transformer, which has strong contextualization capabilities to integrate information from each element in the sequence. A CLS token is added to the top of the sequence to filter out irrelevant cell representations and improve model robustness. The self-attentive mechanism of Transformer is used to fuse the latent representations corresponding to the CLS token, which are then fed into the classifier constructed by a linear layer for classification.

### MID Training and Inference

During the training process, we used the standard supervised training method with cross-entropy as the loss function, along with learning rate warm-up and cosine decay techniques. In order to account for various experimental and computational factors, as well as potential inaccuracies in the labels themselves, we also employed label smoothing to reduce the model’s confidence.

In the testing phase, we made a slight modification to the training approach. While in training we randomly selected 12 cell-slice images to form multiple cell-slices sequences, in testing we increased the number of cell slices included in each sequence. To determine the number of cell slices used in the test phase, we tested the performance of the model with different patch numbers selected. As shown in Figure 1, the overall performance of the model increases with the increase of patch number. In order to take into account the performance of the model and the consumption of the calculation at the same time, we randomly sampled 50 sets of multiple cell slices and computed the average classification result from those sequences as the final outcome. This approach offers the advantage of improving the chances of selecting valuable cell-slice images while also reducing computational costs by not evaluating all individual cells.

### MID attention map

As we utilized the CLS token embeddings generated by Transformer for classification purposes, we obtained the attention map by calculating the dot product between the query representation and key representation among the tokens in Transformer. By applying the Softmax function, each element of the attention map was assigned a value ranging from 0 to 1. Subsequently, we visualized the individual attention map values as solid dots with distinct colors and labeled the corresponding cells in the cell phenotype image (Fig. 3b).

### Cluster and statistical analysis

To evaluate the performance of feature extraction methods for cell-slice image analysis, we employed the t-SNE algorithm in the python sklearn package to reduce the dimensionality of features or embeddings computed by MID, CellProfiler, and DeepProfiler. We then examined the resulting component distributions to investigate the relationship between MID and the cell phenotype observed in images, as well as the effectiveness of MID compared to CellProfiler or DeepProfiler in capturing useful information. To assess the clustering performance, we used the silhouette_score method in sklearn to calculate the silhouette coefficient. For the statistical analysis presented in Figure 5, we applied the Kruskal□Wallis test to test the null hypothesis that the MMP or ROS intensity of images treated with compound at six different concentrations (0.04 μM, 0.12 μM, 0.37 μM, 0.11 μM, 3.33 μM, and 10 μM) were equal, setting significance at p < 0.05.

## Supporting information

https://docs.google.com/spreadsheets/d/1zKDKcuhGKoMO1rk7slskMkrfHhtTj-AH/edit?usp=sharing&ouid=100146808762494663616&rtpof=true&sd=true

https://docs.google.com/document/d/1SzgV5-L7uVfyR9FakrXVlg5rVHygGkep/edit?usp=sharing&ouid=100146808762494663616&rtpof=true&sd=true

## DATA AVAILABILITY

The data used in this article will be provided after the paper is accepted ‘in principle’.

## CODE AVAILABILITY

The MID code was implemented in Python using the deep learning framework of PyTorch. Code, trained models, and scripts reproducing the experiments of this article will be provided after the paper is accepted ‘in principle’.

## AUTHOR INFORMATION

All authors have given approval to the final version of the manuscript.

### Contributions

Xiaodong W and Lipeng L conceived this research. Fan Z, Mengcheng Y and Xueyu G curated the dataset. Xiangrui G, Fan Z and Xueyu G performed data analysis. Xiangrui.G devised deep learning algorithms. Xiaoxiao W ang Dong C conducted the HCS experiments. Xiangrui Gao and Xueyu Guo wrote and modified the paper. Xiaodong W and Lipeng L supervised this work.

## ACKNOWLEDGEMENTS

The authors thank Genwei Zhang for helpful comments and proofreading.

## ABBREVIATIONS

ccRCC: clear-cell renal cell carcinoma
CLS token: Classification Token
CRISPR-cas: Clustered Regularly Interspaced Short Palindromic Repeat-CRISPR-associated (Cas) systems
DMSO: Dimethyl sulfoxide
HCA: High-content analysis
HCS: high-content screening
hEGR: the human Ether-à-go-go-Related Gene
hiPSC-CMs: human induced pluripotent stem cell-derived cardiomyocytes
GT-3 HCV: hepatitis C genotype 3
MID: More Is Different
MMP: mitochondrial membrane potential
MOA: mechanism of action
MPT: Mitochondrial Permeability Transition
OXPHOS: oxidative phosphorylation
RNA: Ribonucleic Acid
ROS: mitochondrial reactive oxygen species
VHL: von Hippel-Lindau

## REFERENCES

1. Gustafsdottir SM, Ljosa V, Sokolnicki KL, Anthony Wilson J, Walpita D, Kemp MM, Petri Seiler K, Carrel HA, Golub TR, Schreiber SL, Clemons PA. Multiplex cytological profiling assay to measure diverse cellular states. PloS one. 2013 Dec 2;8(12):e80999.

2. Stirling DR, Swain-Bowden MJ, Lucas AM, Carpenter AE, Cimini BA, Goodman A. CellProfiler 4: improvements in speed, utility and usability. BMC bioinformatics. 2021 Dec;22(1):1–1.

3. Moshkov N, Bornholdt M, Benoit S, McQuin C, Smith M, Goodman A, Senft R, Han Y, Babadi M, Horvath P, Cimini BA. Learning representations for image-based profiling of perturbations. bioRxiv. 2022 Jan 1.

4. Anderson PW. More is different: broken symmetry and the nature of the hierarchical structure of science. Science. 1972 Aug4;177(4047):393–6

5. Gutman GA, Chandy KG, Adelman JP, Aiyar J, Bayliss DA, Clapham DE, Covarriubias M, Desir GV, Furuichi K, Ganetzky B, Garcia ML. International Union of Pharmacology. XLI. Compendium of voltage-gated ion channels: potassium channels. Pharmacological reviews. 2003 Dec 1;55(4):583–6.

6. Sanguinetti MC, Tristani-Firouzi M. hERG potassium channels and cardiac arrhythmia. Nature. 2006 Mar;440(7083):463–9.

7. Redfern WS, Carlsson L, Davis AS, Lynch WG, MacKenzie IL, Palethorpe S, Siegl PK, Strang I, Sullivan AT, Wallis R, Camm AJ. Relationships between preclinical cardiac electrophysiology, clinical QT interval prolongation and torsade de pointes for a broad range of drugs: evidence for a provisional safety margin in drug development. Cardiovascular research. 2003 Apr 1;58(1):32–45.

8. Haverkamp W, Breithardt G, Camm AJ, Janse MJ, Rosen MR, Antzelevitch C, Escande D, Franz M, Malik M, Moss A, Shah R. The potential for QT prolongation and pro-arrhythmia by non-anti-arrhythmic drugs: clinical and regulatory implications: report on a Policy Conference of the European Society of Cardiology. Cardiovascular research. 2000 Aug 1;47(2):219–33.

9. Fanoe S, Kristensen D, Fink-Jensen A, Jensen HK, Toft E, Nielsen J, Videbech P, Pehrson S, Bundgaard H. Risk of arrhythmia induced by psychotropic medications: a proposal for clinical management. European heart journal. 2014 May 21;35(20):1306–15.

10. Danker T, Möller C. Early identification of hERG liability in drug discovery programs by automated patch clamp. Frontiers in pharmacology. 2014 Sep 2;5:203.

11. Molokanova E, Savchenko A. Bright future of optical assays for ion channel drug discovery. Drug discovery today. 2008 Jan 1;13(1-2):14–22.

12. Deacon M, Singleton D, Szalkai N, Pasieczny R, Peacock C, Price D, Boyd J, Boyd H, Steidl-Nichols JV, Williams C. Early evaluation of compound QT prolongation effects: a predictive 384-well fluorescence polarization binding assay for measuring hERG blockade. Journal of pharmacological and toxicological methods. 2007 May 1;55(3):255–64.

13. Diaz GJ, Daniell K, Leitza ST, Martin RL, Su Z, McDermott JS, Cox BF, Gintant GA. The [3H] dofetilide binding assay is a predictive screening tool for hERG blockade and proarrhythmia: Comparison of intact cell and membrane preparations and effects of altering [K+] o. Journal of pharmacological and toxicological methods. 2004 Nov 1;50(3):187–99.

14. Dykens JA, Will Y. The significance of mitochondrial toxicity testing in drug development. Drug discovery today. 2007 Sep 1;12(17-18):777–85.

15. Begriche K, Massart J, Robin MA, Borgne-Sanchez A, Fromenty B. Drug-induced toxicity on mitochondria and lipid metabolism: mechanistic diversity and deleterious consequences for the liver. Journal of hepatology. 2011 Apr 1;54(4):773–94.

16. Hargreaves IP, Al Shahrani M, Wainwright L, Heales SJ. Drug-induced mitochondrial toxicity. Drug safety. 2016 Jul;39(7):661–74.

17. Tang X, Wang Z, Hu S, Zhou B. Assessing Drug-Induced Mitochondrial Toxicity in Cardiomyocytes: Implications for Preclinical Cardiac Safety Evaluation. Pharmaceutics. 2022 Jun 21;14(7):1313.

18. Soriano A, Miró O, Mensa J. Mitochondrial toxicity associated with linezolid. New England Journal of Medicine. 2005 Nov 24;353(21):2305–6.

19. Meyer JN, Hartman JH, Mello DF. Mitochondrial toxicity. Toxicological Sciences. 2018 Mar 1;162(1):15–23.

20. Bray MA, Gustafsdottir SM, Rohban MH, Singh S, Ljosa V, Sokolnicki KL, Bittker JA, Bodycombe NE, Dančík V, Hasaka TP, Hon CS. A dataset of images and morphological profiles of 30 000 small-molecule treatments using the Cell Painting assay. Gigascience. 2017 Dec;6(12):giw014.

21. Seal S, Carreras-Puigvert J, Trapotsi MA, Yang H, Spjuth O, Bender A. Integrating cell morphology with gene expression and chemical structure to aid mitochondrial toxicity detection. Communications Biology. 2022 Aug 23;5(1):858.

22. Haghighi M, Caicedo JC, Cimini BA, Carpenter AE, Singh S. High-dimensional gene expression and morphology profiles of cells across 28,000 genetic and chemical perturbations. Nature Methods. 2022 Dec;19(12):1550–7.

23. Vaswani A, Shazeer N, Parmar N, Uszkoreit J, Jones L, Gomez AN, Kaiser Ł, Polosukhin I. Attention is all you need. Advances in neural information processing systems. 2017;30.

24. Ghosh MC, Zhang DL, Ollivierre WH, Noguchi A, Springer DA, Linehan WM, Rouault TA. Therapeutic inhibition of HIF-2α reverses polycythemia and pulmonary hypertension in murine models of human diseases. Blood. 2021 May 6;137(18):2509–19.

25. de Wit S, Glen C, de Boer RA, Lang N. Mechanisms shared between cancer, heart failure and targeted anti-cancer therapies. Cardiovascular Research. 2022 Jul 26.

26. Niemann B, Li L, Simm A, Molenda N, Kockskämper J, Boening A, Rohrbach S. Caloric restriction reduces sympathetic activity similar to beta-blockers but conveys additional mitochondrio-protective effects in aged myocardium. Scientific reports. 2021 Jan 21;11(1):1–5.

27. C Pereira G, M Silva A, V Diogo C, S Carvalho F, Monteiro P, J Oliveira P. Drug-induced cardiac mitochondrial toxicity and protection: from doxorubicin to carvedilol. Current pharmaceutical design. 2011 Jul 1;17(20):2113–29.

28. Lozano-Sepúlveda SA, Rincón-Sanchez Ar, Rivas-Estilla AM. Antioxidants benefits in hepatitis C infection in the new DAAs era. Annals of hepatology. 2019 May 1;18(3):410–5.

29. Reshi ML, Su YC, Hong JR. RNA viruses: ROS-mediated cell death. International journal of cell biology. 2014 May 8;2014.

30. Zaninetti C, Gresele P, Bertomoro A, Klersy C, De Candia E, Veneri D, Barozzi S, Fierro T, Alberelli MA, Musella V, Noris P. Eltrombopag for the treatment of inherited thrombocytopenias: a phase II clinical trial. Haematologica. 2020 Mar;105(3):820.

31. Kim TO, Despotovic J, Lambert MP. Eltrombopag for use in children with immune thrombocytopenia. Blood advances. 2018 Feb 27;2(4):454–61.

32. Yang R, Li J, Jin J, Huang M, Yu Z, Xu X, Zhang X, Hou M. Multicentre, randomised phase III study of the efficacy and safety of eltrombopag in Chinese patients with chronic immune thrombocytopenia. British Journal of Haematology. 2017 Jan;176(1):101–10.

33. Ligneau X, Shah RR, BerrebiDBertrand I, Mirams GR, Robert P, Landais L, MaisonDBlanche P, Faivre JF, Lecomte JM, Schwartz JC. Nonclinical cardiovascular safety of pitolisant: comparing International Conference on Harmonization S7B and Comprehensive in vitro ProDarrhythmia Assay initiative studies. British journal of pharmacology. 2017 Dec;174(23):4449–63.

34. Yang X, Ribeiro AJ, Pang L, Strauss DG. Use of human iPSC-CMs in nonclinical regulatory studies for cardiac safety assessment. Toxicological Sciences. 2022 Dec;190(2):117–26.

35. Shinlapawittayatorn K, Chattipakorn S, Chattipakorn N. Effect of sildenafil citrate on the cardiovascular system. Brazilian journal of medical and biological research. 2005;38:1303–11.

