## Supplementary material for "MORE IS DIFFERENT: DRUG PROPERTY ANALYSIS ON CELLULAR HIGH-CONTENT IMAGES USING DEEP LEARNING": https://docs.google.com/document/d/1SzgV5-L7uVfyR9FakrXVlg5rVHygGkep/edit?usp=sharing&ouid=100146808762494663616&rtpof=true&sd=true

Published online: xxx 2023

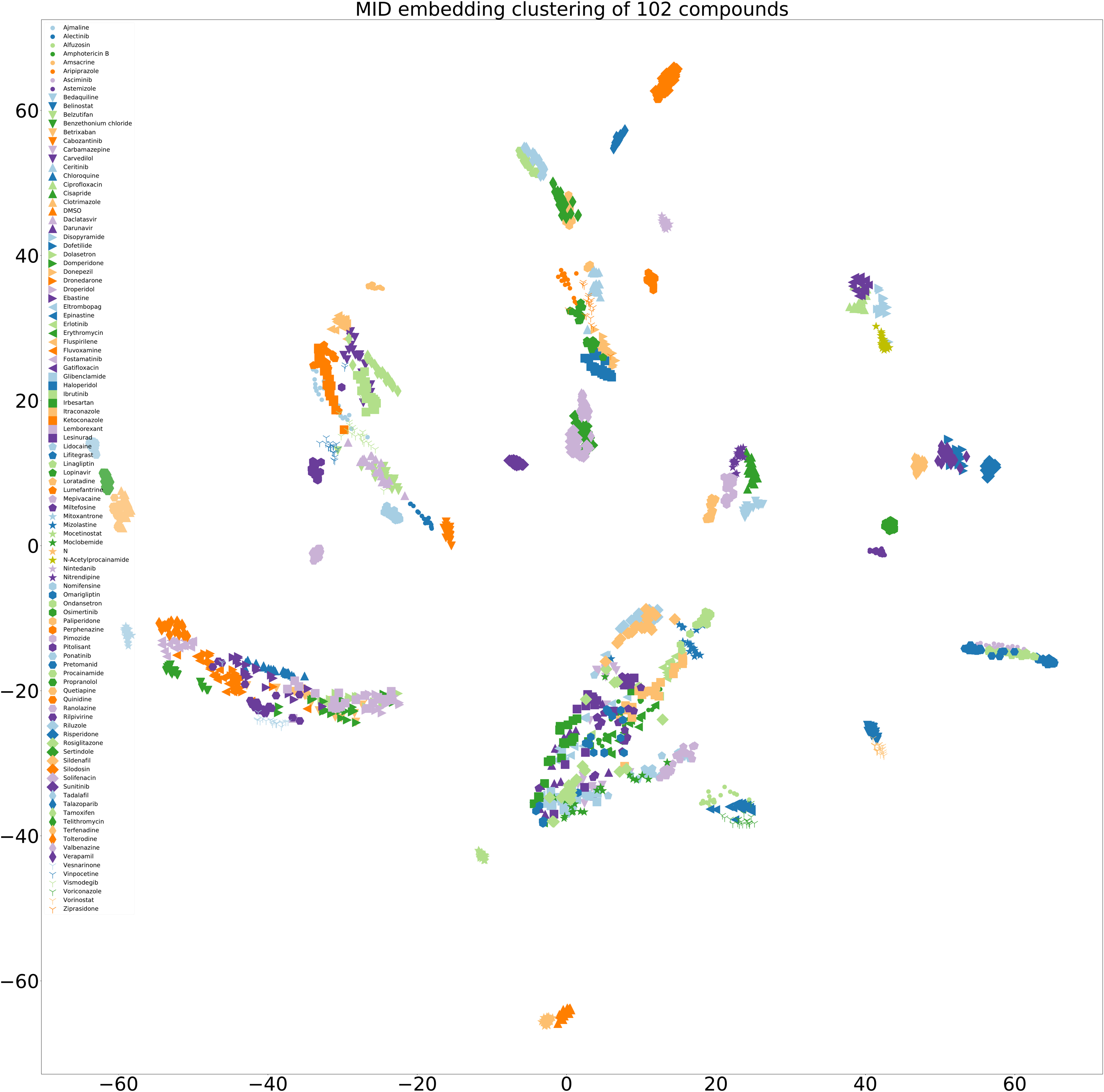
**Supplementary Figure 1: The clusters of MID embeddings of all the 100 compounds, plus DMSO and water (N).**
